# Interferon-Regulatory Factor 7: A Neuroimmune Role for Vapor-Induced Escalations in Ethanol Self-Administration

**DOI:** 10.64898/2026.04.01.715945

**Authors:** Dennis F Lovelock, Joseph M Carew, Elizabeth M McNair, Baylee M Materia, Shadi Z Darawsheh, Anthony M Downs, Sarah E Sizer, Samuel A McDonald, Zoe A McElligott, Leon G Coleman, Joyce Besheer

## Abstract

Neuroimmune signaling is increased in postmortem brain tissue from individuals with alcohol use disorder (AUD), and growing evidence suggests that it contributes to persistent alcohol-related neuroadaptations. Interferon regulatory factor 7 (IRF7), a transcription factor downstream of endosomal Toll-like receptor signaling, is induced in alcohol-relevant brain regions and may contribute to escalated drinking. Here, we tested whether chronic intermittent ethanol (CIE) vapor exposure engages IRF7 signaling during subsequent alcohol self-administration and whether this is associated with altered molecular E/I balance in the aIC and altered functional E/I balance in aIC➔nucleus accumbens projection neurons. Female Wistar rats (n=30) were trained to self-administer alcohol (15% v/v; FR2 vs inactive lever) during 30-minute sessions. After establishing baseline drinking, rats underwent 1-3 cycles of CIE, which increased alcohol self-administration at the 72 h post vapor test. This increase positively correlated with IRF7 levels in the anterior insular cortex (aIC) and nucleus accumbens, while molecular, and immunofluorescence showed that CIE shifted aIC excitatory/inhibitory (E/I) balance toward reduced excitation. Electrophysiological recordings further showed reduced functional E/I balance in aIC neurons projecting to the nucleus accumbens. Knockdown of IRF7 in the aIC attenuated CIE induced escalation of alcohol self-administration, supporting a role for insular IRF7 signaling in alcohol related neuroadaptations that promote escalated drinking.

## INTRODUCTION

Alcohol use disorder (AUD) is a chronic relapsing condition associated with high rates of morbidity and mortality. While many of the acute effects of alcohol on neurological function are well characterized, increasing evidence suggests that chronic alcohol exposure engages complex biological processes that persist beyond intoxication and withdrawal (Heilig et al. 2019). Among these, neuroimmune signaling has emerged as a factor associated with alcohol-related pathology. Postmortem studies of individuals with AUD report elevated expression of immune-related genes and inflammatory markers in the brain (Crews et al. 2013a, 2026; Coleman Jr and Crews 2018), with comparable changes observed in animal models of chronic alcohol exposure (He and Crews 2008; Qin et al. 2008; Crews et al. 2026). Recently, we reported that blocking pro-inflammatory signaling in brain prevents alcohol-induced negative affect suggesting a role for neuroinflammation in AUD pathogenesis (McNair et al. 2026). These findings have prompted a growing interest in neuroimmune mechanisms that shape alcohol-related neuro-adaptations.

Toll-like receptors (TLRs) are innate immune sensors that detect conserved pathogen– and damage-associated molecular patterns in the extracellular and endosomal environment, initiating downstream immune signaling cascades. The expression of several TLRs (e.g., TLR2, 3, 4, and 7) is increased in postmortem human AUD brain (Crews et al. 2013b, 2017; Vetreno et al. 2021). Activation of these receptors induces immune-related gene expression changes in alcohol-relevant brain regions such as the anterior insula (aIC), medial prefrontal cortex (mPFC), and the nucleus accumbens (Acb) (Crews et al. 2013a; Randall et al. 2019; Warden et al. 2019; Lovelock et al. 2022a). Several TLRs have also been implicated in alcohol drinking. For example, repeated activation of TLR3 increases voluntary and operant ethanol intake in rodents (Randall et al. 2019; Warden et al. 2019; Blednov et al. 2021; Lovelock et al. 2022b), and similar increases in drinking are observed following repeated activation of TLR7 (Grantham et al. 2020; Lovelock et al. 2022a). These findings indicate that stimulation of TLRs that are increased in AUD can promote ethanol intake. However, it remains unclear if TLR signaling contributes to the escalation of drinking after high level ethanol exposure seen in AUD and if mediators downstream of TLRs represent therapeutic targets for AUD.

Interferon-regulatory factor 7 (IRF7) is a transcriptional regulator that is engaged downstream of multiple endosomal TLR signaling pathways via activation of the Toll-interleukin receptor (TIR) domain-containing adaptor inducing interferon β (TRIF) signaling to induce type I interferon gene (IFN) expression programs (Honda et al. 2005). TLR7 agonism robustly increases IRF7 expression in regions associated with AUD phenotypes such as the aIC, prefrontal cortex, and Acb (Lovelock et al. 2022a). Critically, in rodent models IRF7 is elevated at the timepoint that corresponds with increased ethanol intake (Grantham et al. 2020; Lovelock et al. 2022a), positioning IRF7 as a potential molecular driver downstream of multiple TLRs that could contribute to neuroadaptations underlying escalated drinking behaviors associated with AUD.

Neuroimmune signaling can influence synaptic plasticity and structure in multiple contexts. This includes altering receptor composition, neurotransmitter release, synaptic spine density and other phenotypes (Mancini et al. 2021; Umpierre and Wu 2021; Zhao et al. 2024). Pro-inflammatory cytokines such as IFNs can alter neurotransmitter receptors, long-term potentiation, neuronal activity and excitatory/inhibitory (E/I) balance (Mendoza-Fernandez et al. 2000; Guo et al. 2016; Mandolesi et al. 2017). E/I balance impacts neuronal circuit function and encoding and disruptions to this balance can result in disease states (Barral and D Reyes 2016; Zhou and Yu 2018). E/I balance can be disrupted by alcohol in different settings (Che et al. 2024). However, the impact of repeated binge ethanol exposure on E/I balance in AUD relevant regions such as the aIC and Acb as well as aIC➔Acb projections, a circuit that is known to regulate drinking, is unknown. Previous imaging studies found reduced aIC➔Acb functional connectivity in young adults with higher alcohol use history, a trend toward a reduction in aIC➔Acb connectivity in rats 24h after chronic binge ethanol (Veer et al. 2019; Crofton et al. 2025). Thus, alterations in aIC➔Acb circuit function might occur after binge ethanol.

The present study was designed to address the role of IRF7 in dependence-induced escalation of drinking. First, we examined the impact of chronic intermittent ethanol vapor (CIE) exposure on ethanol self-administration and its relationship to IRF7 expression in the aIC, mPFC, and Acb. We then assessed the impact of CIE on molecular measures of excitatory/inhibitory (E/I) balance in the aIC and Acb, which led us to measure functional E/I balance using electrophysiology in aIC➔Acb projection neurons. Finally, we tested whether reducing IRF7 expression in the aIC within a short-hairpin RNA (shRNA) viral vector could prevent escalations in ethanol self-administration. Together, these experiments aim to determine whether IRF7 is both engaged by chronic ethanol exposure and functions as a driver of escalated drinking, potentially identifying a target for future therapeutic interventions.

## MATERIALS AND METHODS

### Animals

Adult female Wistar rats (Charles Rivers Laboratories, Wilmington, MA, USA) aged seven weeks upon arrival were used in these experiments. All animals were pair housed in ventilated cages. Rats had ad libitum access to food and water, and the vivarium was on a 12hr light/dark cycle (lights on at 7:00am) and temperature/humidity regulated. All experiments were conducted during the light cycle. Rats were cared for by veterinary staff from the Division of Comparative Medicine at UNC-Chapel Hill. All procedures followed regulations set by the NIH Guide to Care and Use of Laboratory Animals and institutional guidelines.

### Drugs

Ethanol (95% v/v; Pharmco-AAPER, Shelbyville, KY) was diluted in tap water for self-administration sessions, and vaporized (undiluted) for chronic intermittent ethanol exposure.

### Stereotaxic surgery and viral injections

Rats anesthetized via 3% isoflurane and 2% oxygen received a bilateral infusion of a viral vector encoding three short-hairpin RNA (shRNA) sequences (VectorBuilder, Chicago, IL) targeting IRF7 (pAAV8[3miR30]-CAG>EGFP; Vector ID: VB230209-1211tgj, Lot #230310AAVN09), a scrambled control virus (pAAV[miR30]-CMV>EGFP:Scramble_miR30-shRNA:WPRE; Vector ID: VB010000-9397wgw, Lot #221225AAVD16), or aCSF into the aIC (0.8µL/side, AP +2.0, ML ±4.4, D/V –6.0 from skull (Paxinos and Watson 2007) via digital microinjector (Stoelting, Wood Dale, IL) with 1.0µl Hamilton syringes (Hamilton Robotic, NV) and a 32-gauge needle (Stoelting; 0.2µl/min flow rate across 4 min (Ounallah-Saad et al. 2014; Ji and Neugebauer 2019). The injector was left in place for 4 min for virus diffusion.

### Chronic-intermittent ethanol vapor (CIE) exposure

Rats underwent ethanol vapor exposure for 16 hours per day (5 PM to 9 AM) across four consecutive days (Monday PM to Friday AM). Rats were placed in vapor chambers (2 rats per chamber) (La Jolla Alcohol Research, Inc., La Jolla, CA, USA) and ethanol was evaporated into fresh air and dispersed into each cage at a flow rate of ∼16 L/min via an air compressor. Following each daily exposure, tail blood was collected via the tail snip method from two rats per chamber to assess blood alcohol levels (BALs) and adjustments were made to maintain BALs in the 150-220 mg/dl target range. Blood samples were centrifuged, and plasma was analyzed using an AM1 Alcohol Analyzer (Analox Instruments Ltd, Stourbridge, UK). Air controls remained in the home cages in the vivarium.

### EtOH Self-administration

Self-administration sessions (30 min) were conducted in individual operant chambers (Med Associates Inc., St. Albans, VT) enclosed in sound-attenuating boxes equipped with internal fans for air circulation and noise reduction, as previously described (Lovelock et al. 2022a, b). Each chamber had two retractable levers (one on each side) positioned under a cue light and adjacent to a liquid delivery receptacle. Responses on the active lever (fixed-ratio 2; FR2 schedule) delivered 0.1 mL of liquid reinforcer; inactive lever presses had no consequence. Rats were initially trained to self-administer ethanol via a sucrose-fading procedure, starting with 10% (w/v) sucrose + 2% (v/v) ethanol (10S/2E) during an overnight session. Daily self-administration sessions followed with the following progression of solutions: 10S/2E, 10S/5E, 10S/10E, 5S/10E, 5S/15E, 2S/15E, 2S/20E, 20E, and finally 15% ethanol (15E) which was maintained for the remainder of self-administration experiments.

### Tissue Processing for RT-PCR and Immunofluorescence (IF)

At the conclusion of experiments, rats were deeply anesthetized with isoflurane. For RT-PCR studies, rats were perfused with 1X PBS, brains removed and flash frozen, with tissue punches taken from the aIC. For IHC, rats were perfused with PBS followed by 4% paraformaldehyde (PFA). Brains were stored in 4% PFA for 1 week, followed by dehydration in 30% sucrose prior to microtome sectioning at 40µm. Immunofluorescence staining was performed using published protocols (Barnett et al. 2025; McNair et al. 2026). Briefly, free floating tissue sections (40 um) were washed in 0.1 M phosphate-buffered saline (PBS) then incubated for 1 hour at 70°C in pH = 6.0 1x citrate buffer for antigen retrieval (Fisher Scientific, Waltham, MA; NC9935936). To promote membrane permeability, sections were then incubated for 1 hour at room temperature in a blocking solution of 4% normal serum with 0.1% Triton X-100 (Millipore Sigma, Burlington, MA; T8787). Approximately 4 identical sections were stained for each region of interest. Sections were incubated overnight at 4°C in the relevant primary antibody (Table 1) in blocking solution. The next day, sections were washed with PBS and incubated for 1h at room temperature in the appropriate Alexa Fluor-conjugated secondary antibody (1:1000, Invitrogen). Sections were washed, mounted, and cover slips placed with Prolong Gold Anti-Fade mounting medium with DAPI (Thermo Fisher Scientific; P36971). Images were taken on the Keyence BZ-X800 Microscope and analyzed using the accompanying BZ-X800 Analyzer software. Tissue sections were chosen based on desired bregma according to the rat brain atlas (Paxinos and Watson) as follows: infralimbic cortex (IL, +3.00mm), anterior insula (aIC, +3.00mm), nucleus accumbens core (Acb, +2.04mm), and the prelimbic cortex (PrL, +3.00mm). Positive staining was measured as immunoreactive pixels/mm^2^.

**Table 1.** Antibodies for IF.

| Table 1. Antibodies for IF |  |  |  |  |
| --- | --- | --- | --- | --- |
| Target | Isotype | Dilution | Clone | Source; identifier |
| eGFP | Mouse IgG | 1:500 | Monoclonal | Abcam (Cambridge, UK); ab184601 |
| GPHN | Rabbit IgG | 1:100 | Monoclonal | Abcam (Cambridge, UK); ab177154 |
| IRF7 | Rabbit IgG | 1:500 | Polyclonal | Biorbyt (Cambridge, UK); orb6232 |
| PSD95 | Mouse IgG | 1:150 | Monoclonal | Abcam (Cambridge, UK); ab13552 |

### RT PCR

mRNA was extracted from frozen cortex tissue as reported. Samples were homogenized with Trizol (Invitrogen), and RNA was isolated by chloroform extraction. RNA concentration was measured using the nanodrop 2000 ™, and reverse transcription was completed as previously described (Barnett et al. 2025; McNair et al. 2026). SYBR green polymerase chain reaction (PCR) master mix (Applied Biosystems, Foster City, California, USA) was used for quantitative real-time PCR (qRT-PCR) analysis. Primer sequences were designed using the National Library of Medicine Primer-BLAST tool. Primers were only used if they had no predicted non-specific targets and a single peak melt curve (Table 2). Genes of interest were normalized to the expression of housekeeping genes 18S and β-actin using the cycle threshold (Ct) value of each target gene product. The ΔΔCt method was used to compare relative differences between groups, and the percent change relative to the housekeeping gene was used in analysis.

**Table 2.**
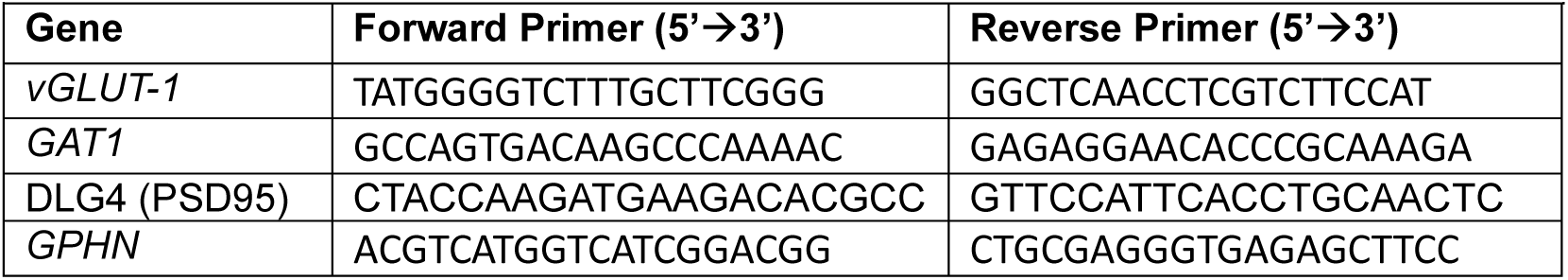
Primers for RT-PCR.

### Electrophysiology

#### Brain slice preparation

Rats were deeply anesthetized with isoflurane, decapitated, and brains were removed and placed into ice-cold sucrose aCSF [in mM: 194 sucrose, 20 NaCl, 4.4 KCl, 2 CaCl2, 1.2 NaH2PO4. 10 glucose, 26 NaHCO3] oxygenated with 95% O2 5% CO2 for slicing. Brains were sliced coronally at 250 µm using a Leica VT1000 vibratome (Germany). Slices containing the aIC were then transferred to a holding container with oxygenated (95% O2 5% CO2) aCSF [in mM: 124 NaCl, 4.4 KCl, 2 CaCl2, 1.2 MgSO4. 1 NaH2PO4, 10 glucose, 26 NaHCO3] at 32 °C for at least 45 minutes. Slices were then transferred to a recording chamber and perfused with oxygenated (95% O2 5% CO2) aCSF [in mM: 124 NaCl, 4.4 KCl, 2 CaCl2, 1.2 MgSO4. 1 NaH2PO4, 10 glucose, 26 NaHCO3] at 30 °C at a constant rate of 2 mL/min.

### Electrophysiological recordings

Electrophysiological recordings were conducted as previously described (Faccidomo et al. 2021; Downs et al. 2023, 2024). Recording electrodes (2-4MΩ) were pulled on a P-97 Micropipette Puller (Sutter Instruments) using borosilicate glass. All voltage clamp recordings were completed using a cesium-methanesulfonate internal solution [in mM: 135 cesium methanesulfonate, 10 KCl, 10 HEPES, 2 MgCl2, 0.2 EGTA, 4 ATP, 0.3 GTP, 20 sodium creatine phosphate] pH ∼7.35, osmolarity ∼285 mOsm. All current clamp recordings were completed using a potassium-gluconate internal solution [in mM: 135 potassium D-gluconate, 5 NaCl, 2 MgCl2, 10 HEPES, 0.6 EGTA, 4 ATP, 0.4 GTP]; pH ∼7.35, osmolarity ∼285 mOsm. All electrophysiology signals were acquired using an Axon Multiclamp 700B and digitized using an Axon Digidata 1550B (Molecular Devices, Sunnyvale, CA). Input resistance, holding current, and access resistance were continuously monitored throughout each experiment, and recordings where access resistance changed by greater than 20% were excluded. 1-3 cells were recorded from each animal for each set of experiments.

Pyramidal neurons projecting from the aIC to the Acb were identified by the presence of a GFP reporter and capacitance > 100 pF. E/I balance was assessed in both spontaneous postsynaptic currents (no TTX) and in miniature postsynaptic currents (500 nM TTX). Neurons were held at –55 mV to isolate glutamatergic excitatory postsynaptic currents (EPSCs) and subsequently held at +10 mV to isolate GABAergic inhibitory postsynaptic currents (IPSCs). The E/I ratio is defined as (EPSC frequency/IPSC frequency), while the synaptic drive ratio is defined as (EPSC Frequency x EPSC amplitude)/(IPSC Frequency x IPSC Amplitude). Resting membrane potential was measured in current clamp at I = 0. Excitability studies were performed in current clamp at I = 0 and with sufficient current injected to hold the cells at –75mV. Rheobase current was determined by applying a series of current ramps (0-120 and 100-220 pA at 120 pA/s). Evoked action potential firing curves were then acquired using step-wise current injections ranging from –100 to +400 pA at 20 pA intervals. Electrophysiology data were analyzed using Easy Electrophysiology (Easy Electrophysiology Ltd., London, UK) or Clampfit (Molecular Devices, San Jose, CA, USA).

### Experiment 1: Self-administration following CIE

Rats (n=30) were trained to self-administer ethanol and established a baseline of drinking behavior over two months. They then underwent three cycles of CIE exposure (n=16 air controls; n=14 CIE), each followed by a 72-hour abstinence period and self-administration sessions on Monday, Wednesday, and Friday to assess post-exposure drinking. A fourth and final cycle of CIE was administered and brains were collected 72-hours post CIE exposure for IHC and/or RT-PCR analyses.

### Experiment 2: IRF7 expression following 1 or 3 cycles of CIE

To determine whether IRF7 elevations were present following fewer cycles of CIE, ethanol naïve rats underwent one week (n=10) or three weeks (n=7) of CIE and brains were collected 72-hours post CIE exposure.

**Experiment 3: (electrophysiology)):** rats (n=12) received pAAVrg-hSyn-EGFP retrograde virus injections into the Acb (A/P: +1.7, M/L: ±1.5, D/V: –6.6 mm), and three weeks later were exposed to one cycle of CIE (6=air controls; 6=vapor). At 72h after the final CIE exposure, brains were collected and slices containing the aIC were prepared. Green, fluorescent cells (i.e., aIC cells projecting to the Acb) were recorded from examining excitatory and inhibitory balance, and excitability. Both spontaneous and miniature (500 nM TTX) EPSCs (–55 mV) and IPSCs (+10 mV) were recorded using Cs-methanosulfonate from each cell. Excitability was measured using K-gluconate intracellular solutions.

### Experiment 4: IRF7 knockdown and self-administration post CIE

To confirm effective knockdown of IRF7 using an shRNA viral vector, rats (n=7) underwent microinjection surgeries where 0.5uL of AAV8.CAG.shIRF7 virus was infused in the right aIC, and scrambled shRNA control virus injected in the left aIC. After 8 weeks to allow for viral expression, brains were collected for IHC analysis of IRF7 levels at the injection site. An additional cohort of rats (N=61) were then trained to self-administer ethanol. Halfway through baseline training, rats underwent bilateral infusions of either AAV8.CAG.shIRF7.eGFP, an AAV8.scrambled RNA control (scRNA) virus, or artificial cerebrospinal fluid (acsf; Tocris Bioscience, Bristol UK). They were then left undisturbed to recover from surgeries for one week. Once recovered, they continued baseline self-administration training and underwent the same CIE and self-administration procedures as described in Experiment 1. Following the fourth cycle of CIE (10-13 weeks post viral injection), brains were collected at 72 hours of abstinence for IHC. To confirm accurate viral injections, eGFP+ expression in the aIC was confirmed using IHC as previously described (AP: +3.00mm to +1.08mm). A total of 8 rats were excluded from the study due to a lack of eGFP+ expression in the aIC (4 Air scCON, 1 CIE scCON, 1 Air shIRF7,and 2 CIE shIRF7).

### Data analysis

Operant self-administration data were measured as the number of ethanol lever responses, and ethanol intake (g/kg) was estimated from body weight and the number of reinforcers delivered. Behavioral data were analyzed using two– or three-way repeated-measures ANOVA as appropriate, BAC, molecular and electrophysiological data were analyzed using unpaired t-tests, and correlations between IRF7 expression and ethanol –related behaviors were assessed using Pearson correlations. All analyses were conducted using GraphPad Prism, with significance set at p < 0.05.

## RESULTS

### CIE promotes escalation of ethanol self-administration and increases IRF7 in aIC, PrL, and Acb

To understand how CIE-induced escalation in drinking affects IRF7 levels, rats were first exposed to three cycles of CIE (or air) with interspersed self-administration. On self-administration days following cycles of CIE, a two-way ANOVA revealed a main effect of CIE on ethanol lever responses [F(1, 28) = 17.07, P < 0.001] and a main effect of session [F(8, 224**) =** 7.825, P < 0.001] (Figure 1B). Post hoc comparisons showed that responding was greater on the first self-administration day post-CIE (Monday) during cycles 1 and 3 compared with subsequent days (Wednesday and Friday; p < 0.05). To further examine this effect, we conducted an additional analysis limited to Mondays (i.e., 72 h after the end of each CIE cycle), which again showed that CIE increased ethanol lever responding relative to the air controls [F(1, 28) = 17.07, p < .001] (Figure 1C) as well as ethanol intake [F(1, 28 = 22.41, p < 0.0001] (**Table 3**). Thus, confirming that CIE produced expected increases in ethanol self-administration especially 72 h after the end of each vapor exposure cycle. We then assessed the impact of CIE on microglial size and expression of the trophic mediator BDNF, which we have reported is reduced in microglia in other AUD-related brain regions (McNair et al. 2026). CIE had no impact on microglia cellular area in the aIC or Acb (**Supplemental Figure 1A-B**) or total BDNF (**Supplemental Figure 1C-D**). However, a reduction in microglial BDNF was seen across both brain regions consistent with a reduction of microglial trophic support to neurons (**Supplemental Figure 1E**). Next, we examined how these dependence-induced drinking escalations are related to IRF7 in the aIC, PrL, IL, and Acb. After the final self-administration session, rats underwent a fourth cycle of CIE (or air), and brains were collected 72 hours after the final ethanol vapor exposure. CIE increased IRF7 in the aIC [t(11) = 2.22, p < 0.05], Acb [t(11) = 6.56, p < 0.0001] and PrL [t(11) = 4.54, p < 0.001] relative to the air controls, with no effect in the IL (Figure 1D-G). IRF7 in the aIC was positively correlated with the percent change in self-administration [r(10) = .60, p < .05] and Acb [r(10) = .69, p < .05] (Figure 1L-M) across groups. However, though CIE increased IRF7 in the PrL and IL, there was no correlation with drinking in these regions (Figure 1N-O). Thus, this suggested a potential functional role of IRF7 in the aIC and Acb on the escalation of drinking. Therefore, we next assessed the impact of CIE on circuit function in the aIC and Acb.

**Figure 1:**
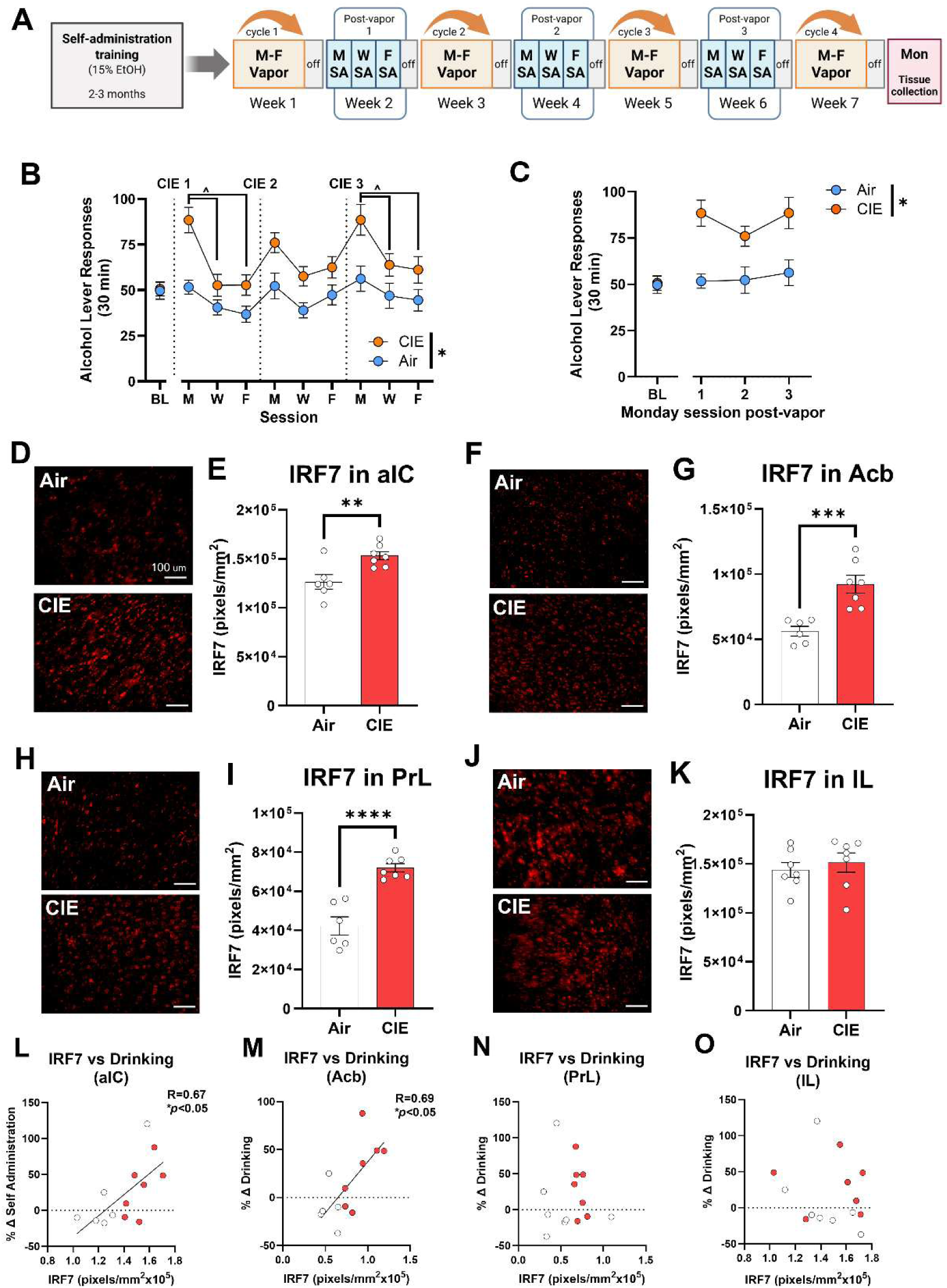
CIE promotes escalation of alcohol self-administration and increases IRF7 expression in alcohol-relevant brain regions. (A) Experiment 1 timeline. Alcohol lever responses at 72 h post-vapor across all sessions (B) and on Mondays only (C). IRF7 immunofluorescence and quantification in the aIC, Acb, PrL, and IL (D-K), and (L-O) correlations between IRF7 immunoreactivity and % increase in drinking (last Monday session vs baseline-BL). *p < 0.05 between CIE and air controls, ^p < 0.05 across post-vapor test days. Scale bar: 100µm

**Table 3:** Ethanol Intake and Inactive Lever Responses for Experiment 1. Table 3. Intake and inactive lever responses from Experiment 1. For intake there was a main effect of CIE [F(1, 28) = 23.73, p<0.0001) and day (F [8, 224] = 8.945, p<0.0001) but no interaction, and there were no effects on inactive lever responses.

| Table 3: Ethanol Intake and Inactive Lever Responses for Experiment 1 |  |  |  |
| --- | --- | --- | --- |
| Measurement | Day | Air | CIE |
| Intake (g/kg) | BL | 0.91 ± 0.06 | 0.95 ± 0.05 |
|  | M1 | 0.93 ± 0.07 | 1.58 ± 0.13 |
|  | W1 | 0.72 ± 0.07 | 0.93 ± 0.10 |
|  | F1 | 0.64 ± 0.07 | 0.95 ± 0.09 |
|  | M2 | 0.87 ± 0.09 | 1.40 ± 0.11 |
|  | W2 | 0.72 ± 0.08 | 1.04 ± 0.08 |
|  | F2 | 0.82 ± 0.09 | 1.11 ± 0.11 |
|  | M3 | 1.00 ± 0.13 | 1.59 ± 0.16 |
|  | W3 | 0.77 ± 0.10 | 1.14 ± 0.10 |
|  | F3 | 0.76 ± 0.08 | 1.10 ± 0.13 |
| Inactive lever responses | BL | 2.92 ± 0.41 | 3.55 ± 0.60 |
|  | M1 | 2.88 ± 0.56 | 1.71 ± 0.47 |
|  | W1 | 2.44 ± 0.56 | 3.36 ± 1.08 |
|  | F1 | 3.38 ± 0.87 | 3.86 ± 1.46 |
|  | M2 | 3.06 ± 0.67 | 2.71 ± 0.66 |
|  | W2 | 3.06 ± 0.64 | 1.57 ± 0.72 |
|  | F2 | 2.31 ± 0.71 | 3.07 ± 0.90 |
|  | M3 | 2.88 ± 0.62 | 3.14 ± 0.74 |
|  | W3 | 4.38 ± 1.01 | 3.93 ± 1.60 |
|  | F3 | 2.44 ± 0.66 | 3.29 ± 0.79 |

### CIE reduces molecular and functional excitatory/inhibitory (E/I) balance in the aIC

E/I balance can be measured globally by assessing GABAergic and glutamatergic signaling by electrophysiology or estimated by molecular markers (Lauterborn et al. 2021). To gain insight into the effect of CIE on aIC and Acb circuit function, we first assessed molecular indices of E/I balance by RT-PCR using presynaptic (vGLUT-1/GAT-1) and post-synaptic (PSD95/GPHN) markers (Lauterborn et al. 2021). CIE had minimal effect on the gene expression of *vGLUT-1*, *GAT1*, or their ratio in the aIC (**Figure 2A-C**) and did not impact the expression of *DLG4*, the gene encoding PSD95 (**Figure 2D**). However, a slight increase in *GPHN* expression was found with a trend toward a significant reduction in the E/I ratio of *DLG4/GPHN* (**Figure 2E-F**). To follow up on this finding, IF analysis of PSD95 and GPHN protein found a 28% reduction in the molecular post-synaptic E/I ratio in the aIC [t(11)=3.038, *p*<0.05; **Figure 2G-H**] that was driven mainly by changes in GPHN (**Supplemental Figure 2A-B**). There were no changes in post-synaptic molecular E/I ratio in the Acb (**Figure 2I**). This suggests there may be disrupted E/I balance of aIC projection neurons. Since aIC➔Acb projections modulate ethanol self-administration (Jaramillo et al. 2018b, a; Haaranen et al. 2020), we next assessed if CIE alters aIC➔Acb projection neurons by electrophysiology.

**Figure 2.**
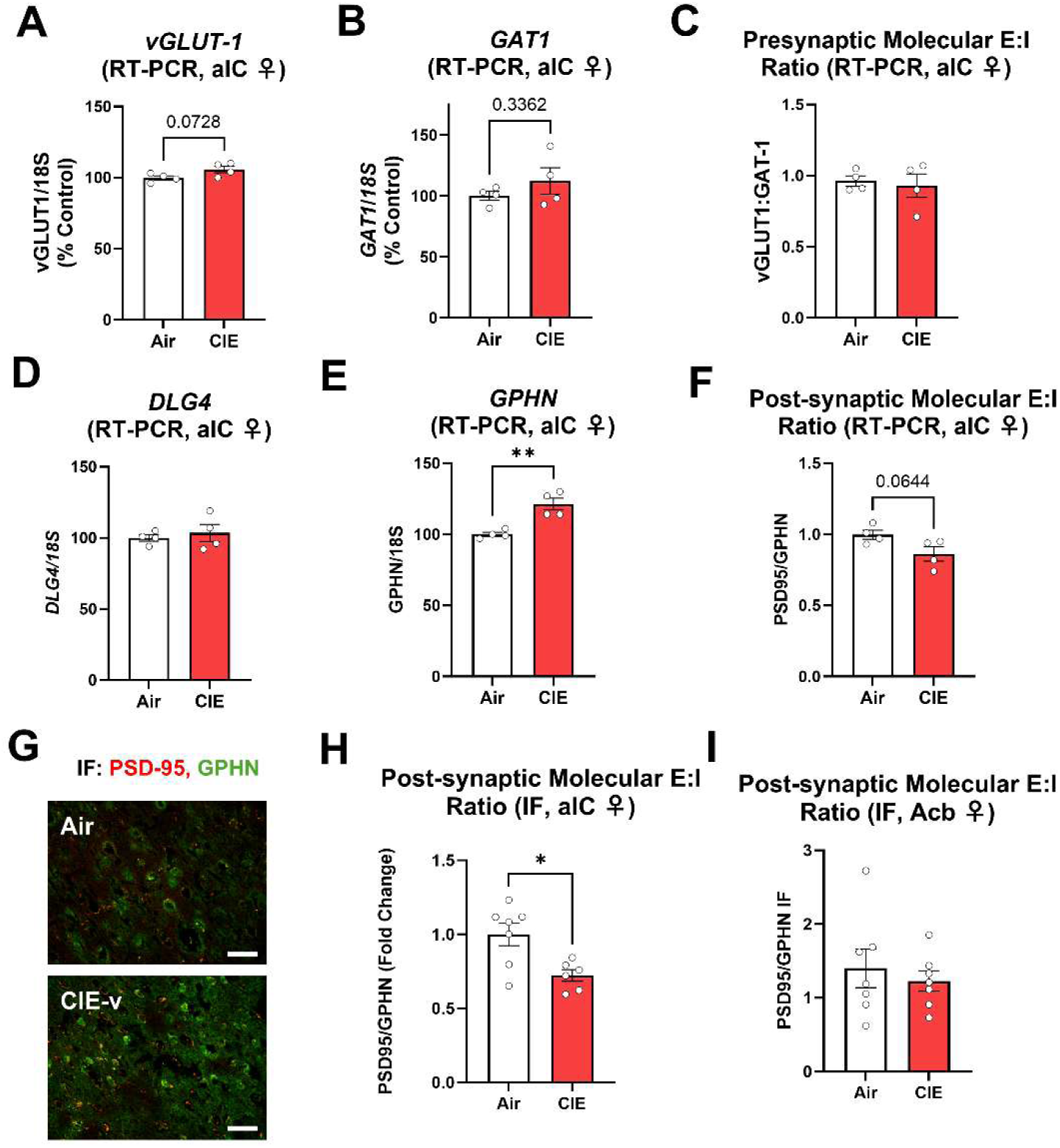
CIE alters molecular E/I marker balance in the anterior insula. RT-PCR in the aIC quantified vGLUT1 and GAT1 (A–B) and their ratio as a presynaptic E/I index (C), DLG4/PSD95 and GPHN (D–E) and their ratio as a postsynaptic molecular E/I index (F). Representative PSD95 and GPHN immunofluorescence in the aIC (G) and the corresponding molecular E/I index are shown for aIC (H) and Acb (I). *p < 0.05 between CIE and air controls. Scale bar: 50µm

The E/I balance of aIC➔Acb projection neurons was evaluated by electrophysiology. One cycle of CIE was used as it was found to produce increased IRF7 [t(8)=6.50, p<0.001] (**Figure 3A-B**) and escalations in ethanol self-administration (**Figure 1B-C**). Rats received stereotaxic injection of retroAAV.eGFP in the Acb and were given three weeks to allow sufficient viral expression. Rats then underwent one cycle of air or CIE with brain slices prepared for electrophysiology 72h after the last ethanol exposure. Electrophysiological recordings were performed on GFP+ aIC projection neurons to the Acb which revealed a significant decrease in E/I balance, consistent with molecular assessment of postsynaptic E/I balance above. Representative tracings illustrate the reduced sEPSC and mEPSC frequency after CIE (**Figure 3C**). When recording aIC➔Acb projection neuron spontaneous postsynaptic currents in the absence of TTX, CIE was found to reduce the E/I ratio [t(12)=3.126, *p<*0.01, 50%] and spontaneous synaptic drive ratio [t(12)=3.835, *p<*0.001, 62%] (**Figure 3D-E**). This was driven primarily by a reduction in the spontaneous excitatory postsynaptic current (sEPSC) frequency [t(12)=5.5, *p<*0.0001, 56%] with a trend toward a reduction in sEPSC amplitude [t(12)=1.93, *p=*0.078] (**Figure 3F-G**). No changes in spontaneous inhibitory postsynaptic current (sIPSC) frequency or amplitude were detected (**Figure 3H-I**). When assessing miniature postsynaptic currents in the presence of TTX, we found similar reductions in E/I balance. CIE reduced both the E/I ratio [t(31)=2.71, *p<*0.05, 50%] and synaptic drive [t(31)=2.5, *p<*0.05, 63%] (**Figure 3J-K**). This was driven primarily by a reduction in mEPSC frequency that showed a trend toward statistical significance [t(31)=1.8, *p=*0.08] (**Figure 3L**). No changes in mEPSC amplitude or inhibitory inputs were found in the presence of TTX (**Figure 3M-O**). We also evaluated the kinetics of both s/mEPSCs and s/mIPSCs. We only identified changes in the decay time of mEPSCs following CIE [t(28)=2.22, p = 0.035), while all other measures of decay and AUC were not significantly different (**Supplemental Figure 3 A-H**). While CIE reduced the E/I balance of aIC➔Acb projections, we found no significant differences in measures of neuronal excitability (action potentials, rheobase, input resistance, resting membrane potential, action potential threshold voltage, or afterhyperpolarization) when cells were held at the common membrane potential of –75 mV or their resting membrane potential (**Supplemental Figure 2C-H**). Thus, CIE reduced the functional E/I balance in aIC➔Acb projection neurons without altering intrinsic neuronal excitability.

**Figure 3.**
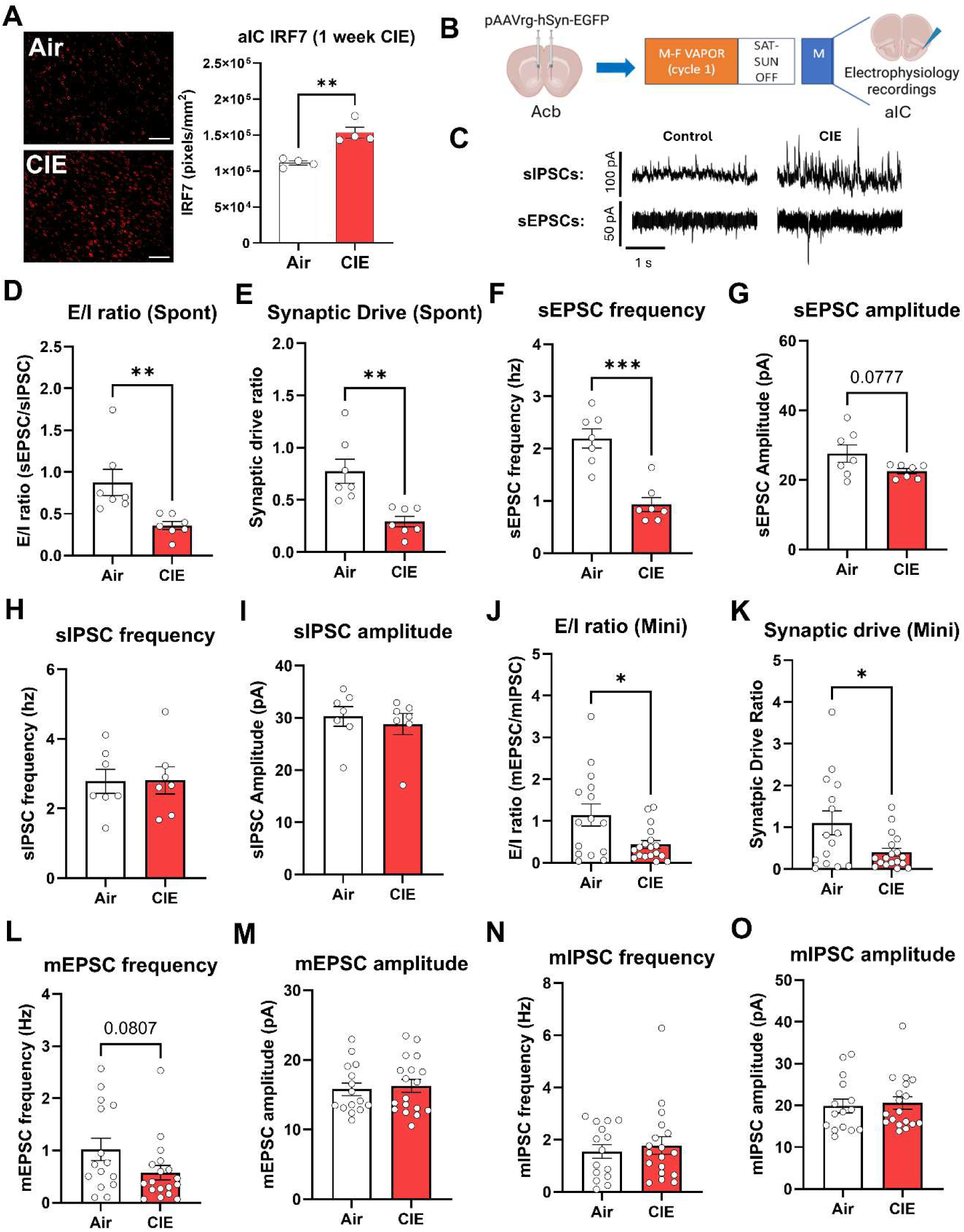
CIE reduces functional E/I balance in aIC➔ Acb projection neurons. (A) Representative IRF7 immunofluorescence in aIC after one cycle of CIE. Scale bar: 100µm (B) Retrograde labeling strategy, CIE timeline, and electrophysiology workflow. (C) Representative traces of sEPSCs and mEPSCs in control and CIE-treated animals. (D-I) Spontaneous E/I ratio, sEPSC and sIPSC frequency and amplitude, and synaptic drive (C–H) Miniature E/I ratio, mEPSC and mIPSC frequency and amplitude, and synaptic drive (J–O). *p < 0.05 between CIE and air controls.

### IRF7 knockdown in the aIC blunts escalation of drinking by CIE

Since aIC IRF7 was strongly correlated with drinking escalation and CIE disrupted E/I balance in the aIC, we next assessed the impact of IRF7 on CIE-induced drinking escalation. To do this, we injected a viral vector containing 3 shRNAs targeting IRF7 (AAV8.CAG.shIRF7[x3]) into the aIC. Since IRF7 is present in both neurons and non-neuronal cells, we chose the ubiquitous CAG promoter for its ability to strongly drive transgene expression across cell types. To verify viral knockdown efficacy, a cohort of alcohol-naïve rats received injection of either scrambled RNA control virus (AAV8.CAG.scRNA), or ethanol virus into the aIC on contralateral hemispheres. Hemispheres that received shIRF7 viral injections showed significantly reduced IRF7 protein near the injection site compared to their within subject contralateral scRNA control virus injection site [t(12)=4.55, p<0.001] (**Supplemental Figure 4A-C**). Next, another cohort of rats wase trained to self-administer ethanol and received bilateral AAV8.CAG.shIRF7[x3].eGFP into the aIC prior to three cycles of CIE or air exposure (**Figure 4A-B**). Separate groups of control rats received either bilateral aCSF or AAV8.CAG.scRNA control virus. Tissue was collected for either for IF or RT-PCR. These control groups did not show differences in ethanol self-administration or IRF7 expression, therefore they were combined. shIRF7 was again found to reduce IRF7 when measured by IF within eGFP+ regions (**Supplemental Figure 4D-E**).On self-administration days following cycles of CIE, a three-way ANOVA revealed a main effect of session on ethanol lever responses [F(8, 344) = 5.92, p < 0.0001] (**Figure 4C**). Post hoc tests indicated that Monday responding was higher than Friday in cycle 1 (p < 0.05) and higher than both Wednesday and Friday in cycle 2 (p < 0.05). By cycle 3, however, this Monday elevation was no longer observed in rats that received IRF7 knockdown.

**Figure 4.**
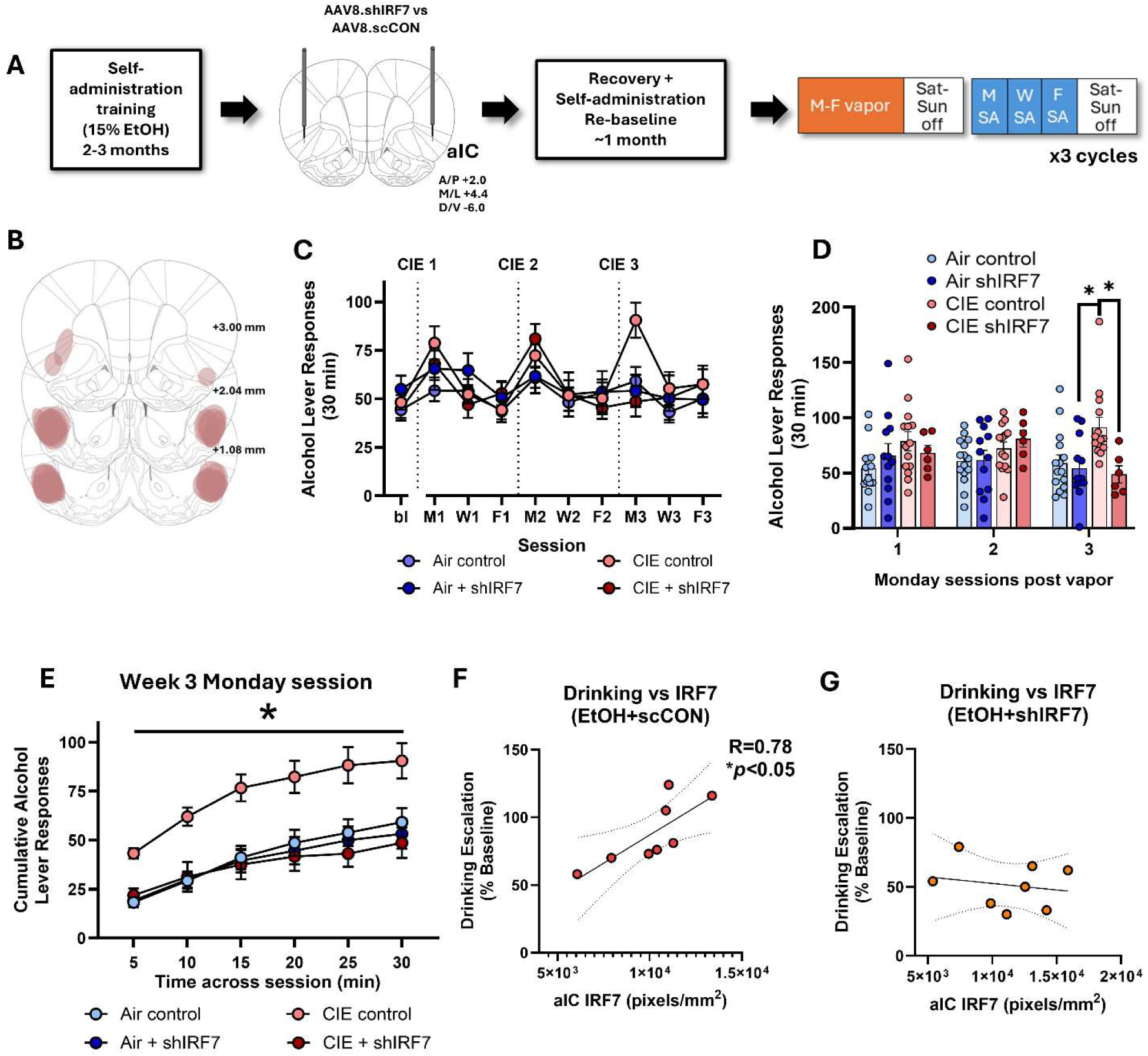
Knockdown of IRF7 in the anterior insula attenuates escalation of alcohol self-administration after CIE. (A) Experimental timeline. (B) Viral injection target and injection sites (C). 72 h post-vapor alcohol lever responses across all sessions (D), on Mondays (E), and across the session on the Monday of week 3 (F). Correlations between drinking escalation and aIC IRF7 immunoreactivity for control (G) and shIRF7 (H) subjects. *p < 0.05.

We next focused on the Monday self-administration sessions to better resolve group differences across CIE cycles. A three-way ANOVA revealed a main effect of CIE on lever responses [F(1, 43) = 4.27, p < 0.05], as well as a week × virus interaction [F(2, 86) = 5.40, p < 0.01] and a three-way interaction [F(2, 86) = 3.10, p < 0.05] (**Figure 4D**). Post hoc tests found that group differences emerged only in week 3, where the CIE control group exhibited greater ethanol lever responding than the Air+shIRF7 and CIE+shIRF7 groups (p < 0.05), indicating that IRF7 knockdown attenuated the CIE-induced escalation in ethanol lever responding. Examination of ethanol intake by three-way ANOVA revealed a significant main effect of group [F(3, 43) = 3.39, p < 0.01] and a significant session × group interaction [F(9, 129) = 3.62, p < 0.001], and, similar to lever responses, post hoc tests showed that CIE controls consumed more ethanol than CIE+shIRF7 rats (p < 0.05) in week 3 (**Table 4**).

**Table 4:**
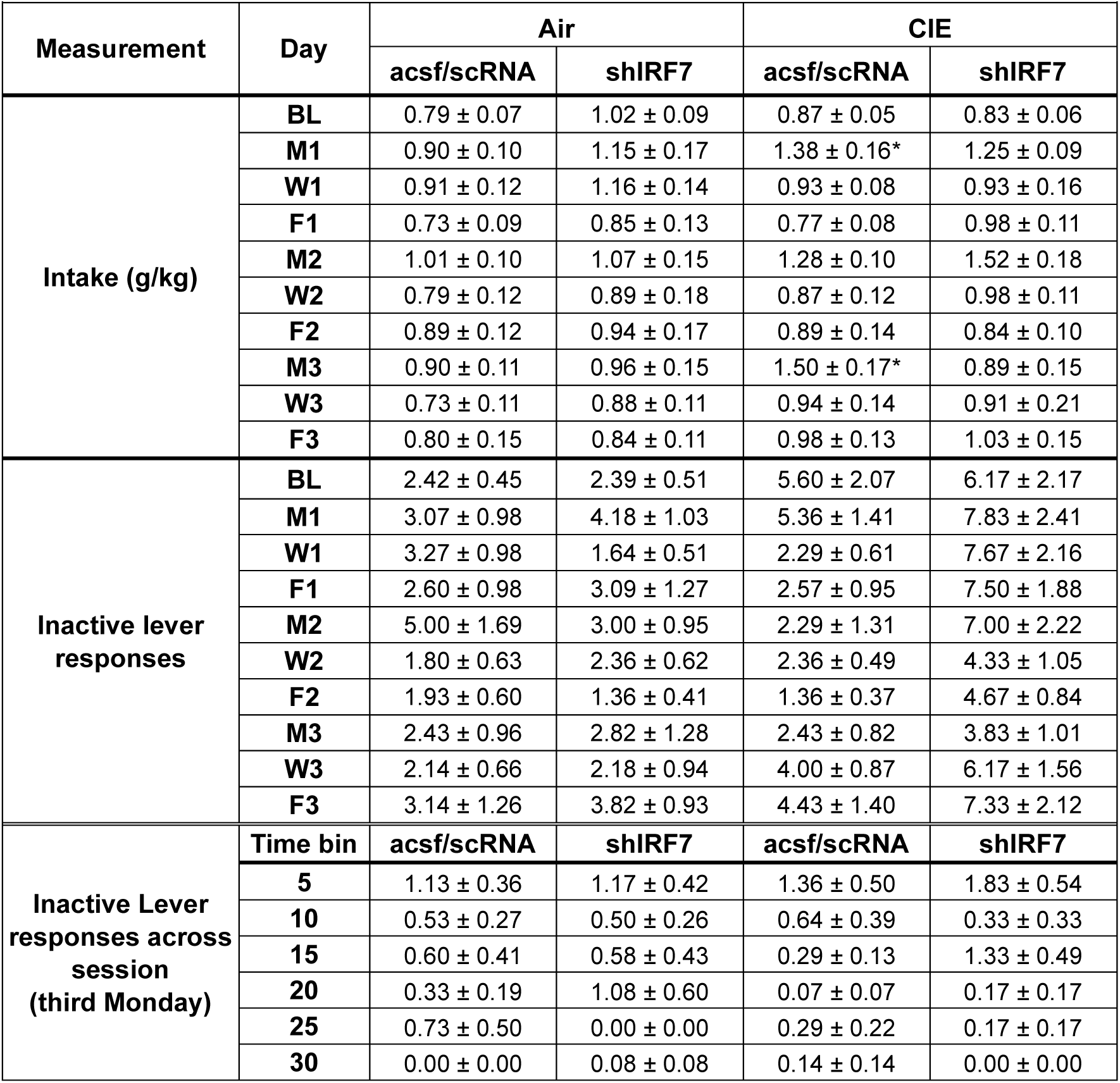
Ethanol Intake and Inactive Lever Responses for Experiment 4. Table 4. Intake and inactive lever responses from Experiment 4. Across days there was a main efect of day [F(8, 328) = 6.40, p<0.0001] and an interaction [F(24, 328) = 1.70, p<0.05]. For inactive lever responding was very low but there were main effects of day [F(8, 344) = 4.09, p<0.0001] and CIE [F(3, 43) = 3.173, p<0.05] but no interaction. For inactive lever responses across the third Monday session there was a main effect of timepoint [F(5, 22) = 8.11, p<0.0001]. *difference from acsf/scRNA control group in post-hoc test, p<0.05.

Given that group differences were specific to the Week 3 Monday session, we next examined the pattern of ethanol self-administration across that session in 5-min bins. A three-way ANOVA revealed main effects of shIRF7 virus [F(1, 44) = 8.84, p < 0.01], CIE [F(1, 44) = 5.32, p < 0.05], and time bin [F(5, 220) = 73.08, p < 0.0001] (**Figure 4E**). Significant interactions were observed for virus × CIE [F(1, 44) = 6.88, p < 0.05] and bin × virus [F(5, 220) = 2.96, p < 0.05]. Post hoc comparisons indicated that all groups had significantly lower cumulative ethanol lever responses relative to the CIE control group (p < 0.01). Thus, shIRF7 expression attenuated the escalation of self-administration observed in CIE controls, reducing self-administration to levels comparable with non-CIE groups. Overall, these findings indicate that knockdown of IRF7 following CIE was effective at reducing the CIE-induced escalation in ethanol self-administration. Further, aIC IRF7 was positively correlated with drinking after CIE (**Figures 4F**) as above (**Figures 1L**). However, this association was not present in rats that underwent IRF7 knockdown (**Figure 4G**), further supporting a role for IRF7 in CIE escalation of drinking. We also measured molecular markers of post-synaptic E/I ratio in remaining aIC sections. CIE again caused a reduction in molecular post-synaptic E/I ratio that was not present after IRF7 KD (**Supplemental Figure 4F).** This was driven primarily by a reduction in PSD95 with CIE with no changes in GPHN (**Supplemental Figure 4G-J**). Thus, IRF7 KD in the aIC blunts CIE-induced increases in drinking and molecular E/I balance.

## DISCUSSION

In the present study, chronic intermittent ethanol vapor exposure increased operant ethanol self-administration, with the largest increases during the first exposure sessions, 72 hours post vapor. This was accompanied by increased IRF7 immunoreactivity in the PrL, aIC, and Acb, with positive correlations between drinking escalation and IRF7 in the aIC and Acb. Molecular markers supported a shift toward reduced E/I balance in aIC but not Acb, pointing to an insular cortex-anchored shift toward relatively reduced excitation post-CIE. This was confirmed functionally, with electrophysiological recordings finding a >50% reduction in E/I balance of aIC➔Acb projection neurons post-CIE. This was driven primarily by fewer excitatory synaptic events, with inhibitory inputs and intrinsic excitability unchanged, providing a circuit level correlate of the synaptic changes in the aIC. Reducing IRF7 in the aIC attenuated the emergence of escalated ethanol responding across cycles and normalized molecular markers of E/I balance, supporting a functional role for IRF7 signaling in aIC in dependence-induced escalation of drinking.

A critical feature of the present design is the assessment of self-administration at 72h after the last vapor exposure. Notably, escalated drinking and increased IRF7 were evident at 72 h post-CIE after one or multiple CIE cycles, consistent with prior work implicating TLR-related neuroimmune signaling in excessive alcohol drinking and IRF7 induction (Lovelock et al. 2022b, a), and the present study identified concomitant alterations in E/I balance. This interval corresponds to early abstinence, a phase characterized by emerging negative affective and motivational disturbances that can promote escalated alcohol seeking via negative reinforcement mechanisms (Koob and Le Moal 2008; Heilig et al. 2019; Koob 2021). At this timepoint, CIE increased ethanol self-administration, and the magnitude of this increase was positively correlated with IRF7 immunoreactivity in the aIC and Acb. Neuroimmune signaling remains elevated into early abstinence and has been proposed to contribute to the development of these aversive internal states, thereby biasing behavior toward alcohol seeking (Crews et al. 2017; McNair et al. 2026). In line with this framework, the IRF7-related escalation in drinking seen here is consistent with increases in drinking seen after repeated TLR7 or TLR3 stimulation, both of which induce IRF7 (Randall et al. 2019; Warden et al. 2019; Grantham et al. 2020; Lovelock et al. 2022b, a). Further, it indicates that not only does IRF7 induction enhance drinking, but that IRF7 inhibition prevents enhanced drinking after binge exposure. Thus, these studies implicate IRF7 is a potential therapeutic target for AUD.

Innate immune activation is known to influence excitatory synaptic transmission and plasticity (Yirmiya and Goshen 2011; Vezzani and Viviani 2015). We previously reported that repeated TLR3 activation alters glutamatergic gene expression alongside increased ethanol self-administration (Vetreno et al. 2021; Lovelock et al. 2022b). Consistent with this framework, CIE produced a shift toward reduced excitatory influence within the anterior insula, reflected by molecular changes in postsynaptic E/I markers and a functional reduction in spontaneous and miniature excitatory events in aIC➔Acb projection neurons. This was expressed as fewer excitatory synaptic events without changes in inhibitory input or intrinsic excitability, pointing to a synaptic input level adaptation onto aIC➔Acb neurons at this 72h timepoint that coincides with increased drinking that seem consistent with imaging studies (Veer et al. 2019; Crofton et al. 2025). A current question is whether reduced excitatory synaptic input in this projection biases downstream processing to promote escalated drinking. The behavioral consequence of reduced aIC input likely depends on which Acb cell populations are most strongly engaged by this projection. For instance, increases in Acb D1 neuronal activity and reductions in D2 neuronal activity both promote drinking (Strong et al. 2020). Reduced excitatory input on D2 neurons from the aIC could thus promote ethanol intake after dependence. Consistent with this idea, prior work found that reducing ventral hippocampal inputs to the Acb increases ethanol consumption (Griffin et al. 2023). Alternatively, reduced excitatory synaptic events could reflect compensatory synaptic remodeling of aIC➔Acb inputs after alcohol vapor, given that CIE alters glutamatergic plasticity and spine structure in Acb (Jeanes et al. 2011; Lovinger and Kash 2015; Uys et al. 2016). Future studies will dissect the specific aIC➔Acb projections that promote drinking. A limitation of the present work is that electrophysiology recordings were conducted in rats exposed to CIE that did not have a history of operant self-administration. Given that chronic ethanol exposure can alter cortico-striatal synaptic plasticity (Lovinger and Kash 2015) and that cortico-striatal LTP and LTD can bidirectionally control operant ethanol seeking (Ma et al. 2018), prior alcohol self-administration could potentially influence the observed reduction in excitatory synaptic input. This will be important to test in future studies combining ethanol self-administration history with circuit-specific electrophysiology. Notably, these experiments focused on female rats, reflecting their faster escalation trajectories and larger ethanol-induced IRF7 responses reported previously (Lovelock et al. 2022a). Determining whether comparable IRF7-associated circuit adaptations emerge in males in future work will also be important.

Reduced glutamatergic input in aIC➔Acb projection neurons at 72 h post-CIE, together with the positive correlation between IRF7 expression and escalated drinking, motivated direct manipulation of insular cortex IRF7 signaling. Viral knockdown of IRF7 within the aIC attenuated the emergence of escalated ethanol self-administration across repeated CIE cycles. This provides functional support for a role of insular cortex IRF7 signaling in escalated drinking. Notably, increased drinking was evident after a single CIE cycle in both control and shIRF7 groups with the shIRF7 knockdown attenuating the escalation in drinking after the third CIE cycle. This pattern is consistent with a role for insular cortex IRF7 signaling in the progression or maintenance of escalated drinking across repeated CIE cycles. IRF7 expression was elevated and correlated with drinking in both the aIC and Acb, whereas electrophysiological recordings were limited to aIC➔Acb projection neurons. It therefore remains unknown whether functional adaptations within the Acb, or within other afferent regions converging on the Acb, also contribute to the escalated drinking.

Together, these findings support a model in which chronic intermittent ethanol exposure induces IRF7 in anterior insular cortex and nucleus accumbens and shifts excitatory drive within the insular cortex to accumbens pathway during escalated ethanol self-administration. By linking IRF7 expression to both circuit-level E/I changes and the maintenance of escalation across CIE cycles, the present work highlights insular cortex IRF7 as a plausible contributor to dependence-related molecular adaptations that drive escalated drinking. Future studies that manipulate defined insula projections and cell types will be important for clarifying how IRF7-dependent neuroimmune activation shapes cortico-striatal function and alcohol-related behavior.

## Funding

P60: 5P60AA011605-29

INIA U01: U01AA02091

K01: K01DA061050

## References

1. Barnett AM, McNair EM, Dawkins L, et al (2025) Loss of lysosomal acid lipase contributes to Alzheimer’s disease pathology and cognitive decline. Alzheimer’s & Dementia 21:e70486

2. Barral J, D Reyes A (2016) Synaptic scaling rule preserves excitatory–inhibitory balance and salient neuronal network dynamics. Nature neuroscience 19:1690–1696

3. Blednov YA, Da Costa A, Mayfield J, et al (2021) Deletion of Tlr3 reduces acute tolerance to alcohol and alcohol consumption in the intermittent access procedure in male mice. Addiction biology 26:e12932

4. Che C, Zhou T, Peng S-Y, Peng Y-M (2024) Alcohol exposure induces cortical activity change during quiescent state. Neuroscience Letters 843:138012

5. Coleman Jr LG, Crews FT (2018) Innate immune signaling and alcohol use disorders. In: The neuropharmacology of alcohol. Springer, pp 369–396

6. Crews FT, Lawrimore CJ, Walter TJ, Coleman Jr LG (2017) The role of neuroimmune signaling in alcoholism. Neuropharmacology 122:56–73

7. Crews FT, Qin L, Coleman L, et al (2026) Cortical reactive microglia activate astrocytes, increasing neurodegeneration in human alcohol use disorder. Brain Behav Immun 131:106156. 10.1016/j.bbi.2025.106156

8. Crews FT, Qin L, Sheedy D, et al (2013a) High mobility group box 1/Toll-like receptor danger signaling increases brain neuroimmune activation in alcohol dependence. Biological psychiatry 73:602–612

9. Crews FT, Qin L, Sheedy D, et al (2013b) High mobility group box 1/Toll-like receptor danger signaling increases brain neuroimmune activation in alcohol dependence. Biological psychiatry 73:602–612

10. Crofton EJ, Lee S-H, Ban W, et al (2025) Chronic ethanol exposure reduces resting state functional connectivity and regional synchrony in male rats. Psychopharmacology 1–13

11. Downs AM, Catavero CM, Kasten MR, McElligott ZA (2023) Tauopathy and alcohol consumption interact to alter locus coeruleus excitatory transmission and excitability in male and female mice. Alcohol 107:97–107

12. Downs AM, Kmiec G, McElligott ZA (2024) Oral fentanyl consumption and withdrawal impairs fear extinction learning and enhances basolateral amygdala principal neuron excitatory-inhibitory balance in male and female mice. Addiction neuroscience 13:100182

13. Faccidomo S, Cogan ES, Hon OJ, et al (2021) Calcium-permeable AMPA receptor activity and GluA1 trafficking in the basolateral amygdala regulate operant alcohol self-administration. Addiction biology 26:e13049

14. Grantham E, Warden A, McCarthy G, et al (2020) Role of toll-like receptor 7 (TLR7) in voluntary alcohol consumption. Brain, behavior, and immunity 89:423–432

15. Griffin WC, Lopez MF, Woodward JJ, Becker HC (2023) Alcohol dependence and the ventral hippocampal influence on alcohol drinking in male mice. Alcohol 106:44–54

16. Guo J, Zhang W, Zhang L, et al (2016) Probable involvement of p11 with interferon alpha induced depression. Scientific reports 6:17029

17. Haaranen M, Schäfer A, Järvi V, Hyytiä P (2020) Chemogenetic stimulation and silencing of the insula, amygdala, nucleus accumbens, and their connections differentially modulate alcohol drinking in rats. Frontiers in behavioral neuroscience 14:580849

18. He J, Crews FT (2008) Increased MCP-1 and microglia in various regions of the human alcoholic brain. Experimental neurology 210:349–358

19. Heilig M, Augier E, Pfarr S, Sommer WH (2019) Developing neuroscience-based treatments for alcohol addiction: A matter of choice? Translational psychiatry 9:255

20. Honda K, Yanai H, Negishi H, et al (2005) IRF-7 is the master regulator of type-I interferon-dependent immune responses. Nature 434:772–777

21. Jaramillo AA, Randall PA, Stewart S, et al (2018a) Functional role for cortical-striatal circuitry in modulating alcohol self-administration. Neuropharmacology 130:42–53. 10.1016/j.neuropharm.2017.11.035

22. Jaramillo AA, Van Voorhies K, Randall PA, Besheer J (2018b) Silencing the insular-striatal circuit decreases alcohol self-administration and increases sensitivity to alcohol. Behavioural brain research 348:74–81

23. Jeanes ZM, Buske TR, Morrisett RA (2011) In vivo chronic intermittent ethanol exposure reverses the polarity of synaptic plasticity in the nucleus accumbens shell. The Journal of pharmacology and experimental therapeutics 336:155–164

24. Ji G, Neugebauer V (2019) Contribution of corticotropin-releasing factor receptor 1 (CRF1) to serotonin receptor 5-HT2CR function in amygdala neurons in a neuropathic pain model. International Journal of Molecular Sciences 20:4380

25. Koob GF (2021) Drug addiction: hyperkatifeia/negative reinforcement as a framework for medications development. Pharmacological reviews 73:163–201

26. Koob GF, Le Moal M (2008) Neurobiological mechanisms for opponent motivational processes in addiction. Philosophical Transactions of the Royal Society B: Biological Sciences 363:3113–3123

27. Lauterborn JC, Scaduto P, Cox CD, et al (2021) Increased excitatory to inhibitory synaptic ratio in parietal cortex samples from individuals with Alzheimer’s disease. Nature communications 12:2603

28. Lovelock DF, Liu W, Langston SE, et al (2022a) The Toll-like receptor 7 agonist imiquimod increases ethanol self-administration and induces expression of Toll-like receptor related genes. Addiction Biology 27:e13176

29. Lovelock DF, Randall PA, Van Voorhies K, et al (2022b) Increased alcohol self-administration following repeated Toll-like receptor 3 agonist treatment in male and female rats. Pharmacology Biochemistry and Behavior 216:173379

30. Lovinger DM, Kash TL (2015) Mechanisms of neuroplasticity and ethanol’s effects on plasticity in the striatum and bed nucleus of the stria terminalis. Alcohol research: current reviews 37:109

31. Ma T, Cheng Y, Roltsch Hellard E, et al (2018) Bidirectional and long-lasting control of alcohol-seeking behavior by corticostriatal LTP and LTD. Nature neuroscience 21:373–383

32. Mancini A, Ghiglieri V, Parnetti L, et al (2021) Neuro-immune cross-talk in the striatum: from basal ganglia physiology to circuit dysfunction. Frontiers in immunology 12:644294

33. Mandolesi G, Bullitta S, Fresegna D, et al (2017) Interferon-γ causes mood abnormalities by altering cannabinoid CB1 receptor function in the mouse striatum. Neurobiology of disease 108:45–53

34. McNair EM, Dawkins LW, Materia B, et al (2026) Microglia promote neurodegeneration and hyperkatifeia during withdrawal and abstinence from binge alcohol. The American Journal of Pathology 196:306–325

35. Mendoza-Fernandez V, Andrew RD, Barajas-Lopez C (2000) Interferon-α inhibits long-term potentiation and unmasks a long-term depression in the rat hippocampus. Brain research 885:14–24

36. Ounallah-Saad H, Sharma V, Edry E, Rosenblum K (2014) Genetic or pharmacological reduction of PERK enhances cortical-dependent taste learning. Journal of Neuroscience 34:14624–14632

37. Paxinos G, Watson C (2007) The Rat Brain in Stereotaxic Coordinates, 6th Edition. Academic Press, San Diego

38. Qin L, He J, Hanes RN, et al (2008) Increased systemic and brain cytokine production and neuroinflammation by endotoxin following ethanol treatment. Journal of neuroinflammation 5:10

39. Randall PA, Vetreno RP, Makhijani VH, et al (2019) The toll-like receptor 3 agonist poly (I: C) induces rapid and lasting changes in gene expression related to glutamatergic function and increases ethanol self-administration in rats. Alcoholism: Clinical and Experimental Research 43:48–60

40. Strong C, Hagarty D, Brea Guerrero A, et al (2020) Chemogenetic selective manipulation of nucleus accumbens medium spiny neurons bidirectionally controls alcohol intake in male and female rats. Scientific Reports 10:19178

41. Umpierre AD, Wu L-J (2021) How microglia sense and regulate neuronal activity. Glia 69:1637–1653

42. Uys JD, McGuier NS, Gass JT, et al (2016) Chronic intermittent ethanol exposure and withdrawal leads to adaptations in nucleus accumbens core postsynaptic density proteome and dendritic spines. Addiction biology 21:560–574

43. Veer IM, Jetzschmann P, Garbusow M, et al (2019) Nucleus accumbens connectivity at rest is associated with alcohol consumption in young male adults. European Neuropsychopharmacology 29:1476–1485

44. Vetreno RP, Qin L, Coleman Jr LG, Crews FT (2021) Increased Toll-like Receptor-MyD88-NFκB-Proinflammatory neuroimmune signaling in the orbitofrontal cortex of humans with alcohol use disorder. Alcoholism: Clinical and Experimental Research 45:1747–1761

45. Warden AS, Azzam M, DaCosta A, et al (2019) Toll-like receptor 3 dynamics in female C57BL/6J mice: Regulation of alcohol intake. Brain Behav Immun 77:66–76. 10.1016/j.bbi.2018.12.006

46. Zhao S, Umpierre AD, Wu L-J (2024) Tuning neural circuits and behaviors by microglia in the adult brain. Trends in neurosciences 47:181–194

47. Zhou S, Yu Y (2018) Synaptic EI balance underlies efficient neural coding. Front Neurosci 12: 46. Case Sex Age (years) Cause of death Postmortem delay (h) EC layer No AS No SS No all synapses No synapses/μm3 (mean\pmSD) EC layer Type of synapse EC layer Type of synapse Total synapses AS 83:12–9

