## Supplement for "Interferon-Regulatory Factor 7: A Neuroimmune Role for Vapor-Induced Escalations in Ethanol Self-Administration"

**Supplemental Figures**

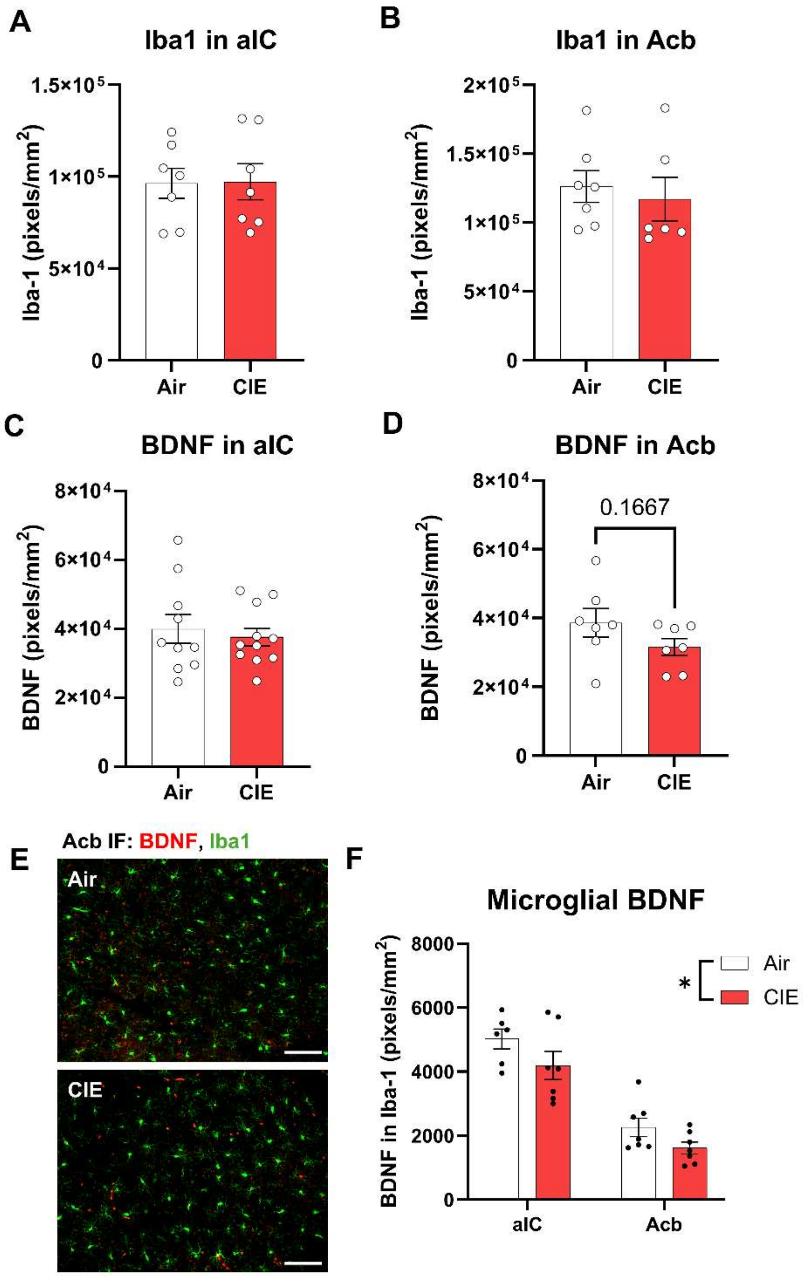


**Supplemental Figure 1. CIE reduces microglial BDNF.** CIE did not alter gross microglial area in the (A) aIC or (B) Acb. (C) CIE did not alter total BDNF in the aIC. (D) CIE caused a trend toward reduced BDNF in the Acb. (E) Representative image of BDNF and Iba1 immunofluorescence in the Acb. Scale bar: 100µm (F) CIE caused a significant reduction in microglial BDNF across the aIC and Acb. $F_{1,23}=5.247$, $p<0.05$.

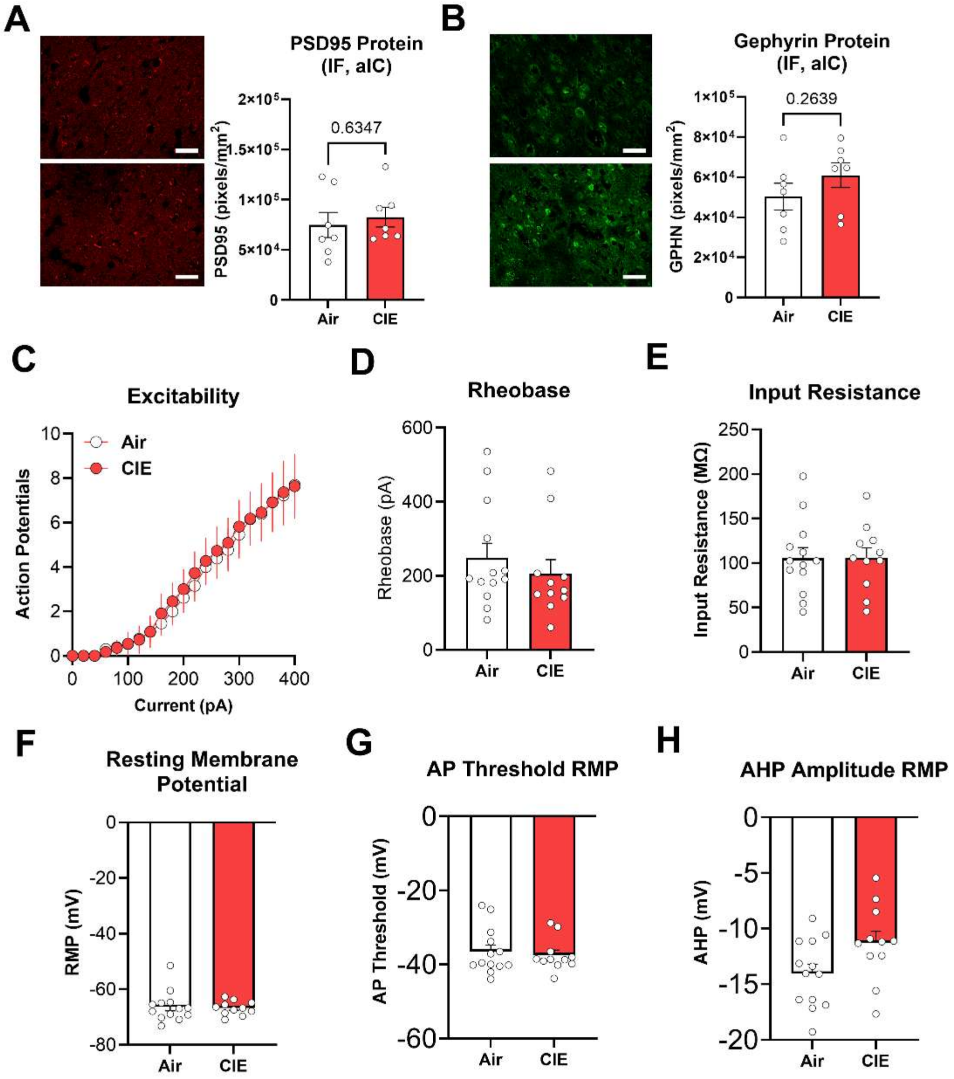


**Supplemental Figure 2. CIE impacts on molecular measures of E/I balance and neuronal excitability.** (A) CIE had no effect on PSD95 in the aIC. (B) CIE caused a trend toward an increase in GPHN. Scale bar: 50μm. (C-H) CIE had no impact on electrophysiological measures of intrinsic excitability including (C) evoked action potential firing, (D) rheobase, (E) membrane input resistance, (F) resting membrane potential, (G) action potential threshold voltage, or (H) amplitude.

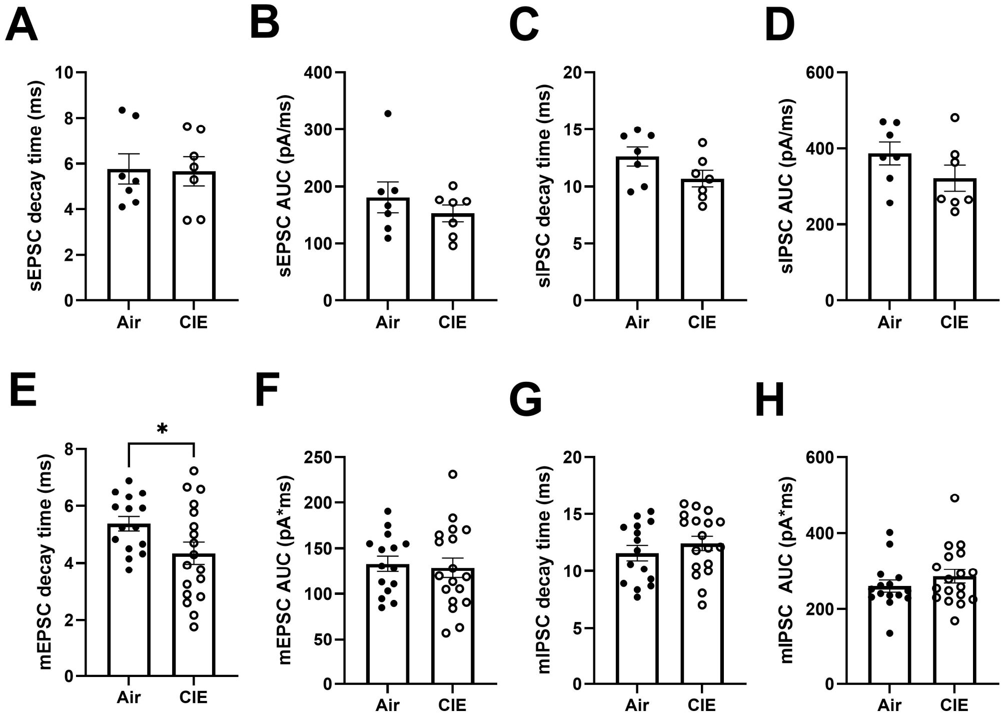


**Supplemental Figure 3. CIE impacts on s/mEPSC and s/mIPSC kinetics.** CIE had no effect on sEPSC decay time (A), sEPSC AUC (B), sIPSC decay time (C), or sIPSC AUC (D). CIE resulted in a signficant decrease in mEPSC decay time (E). There were no signficant changes in mEPSC AUC (F), mIPSC decay time (G), or mIPSC AUC (H).

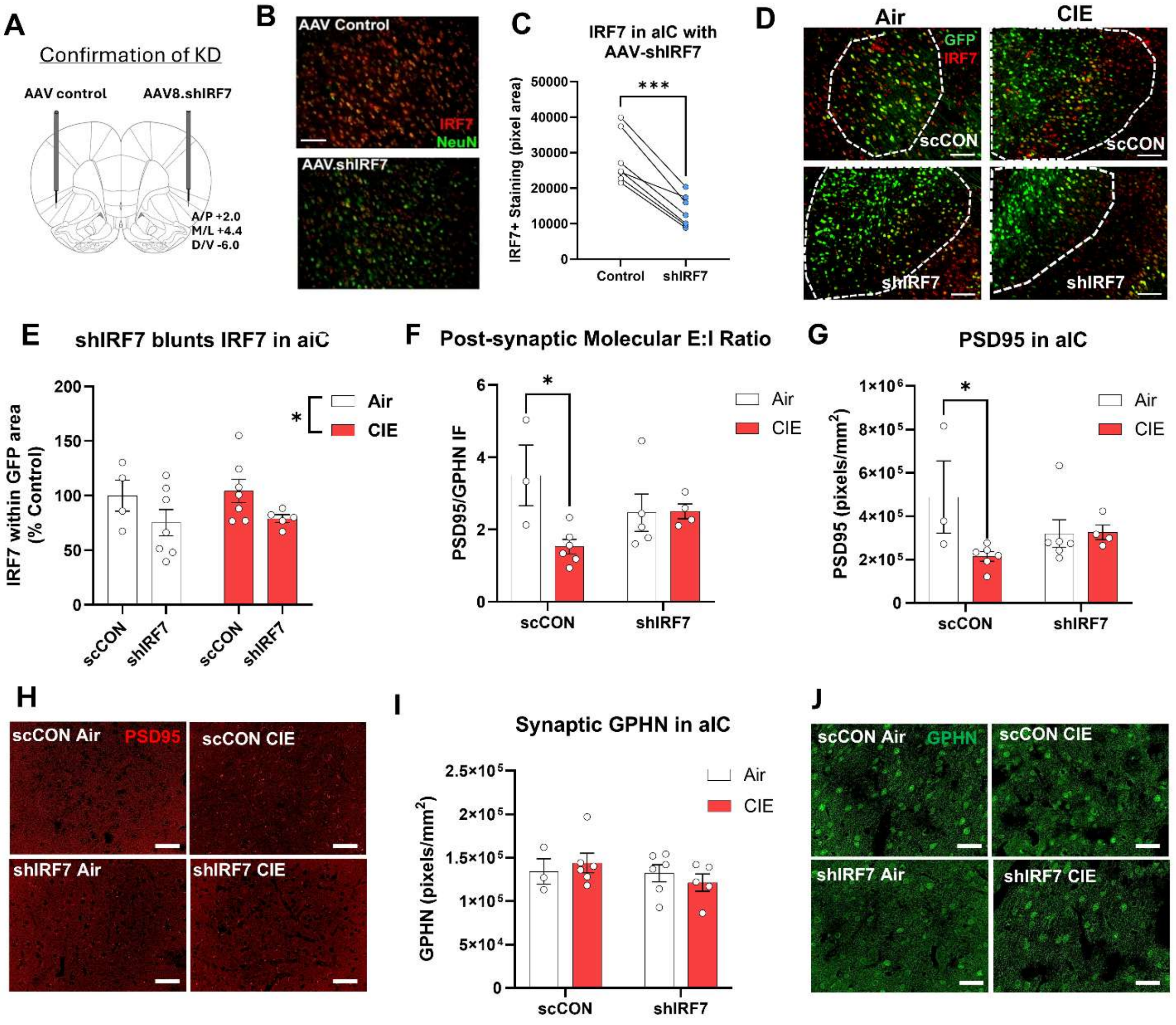


**Supplemental Figure 4. IRF7 knockdown efficiency and impact on molecular E/I markers.** (A) schematic of AAV8.shIRF7 or AAV8.scCON injection into contralateral hemispheres within the same animals. (B) Representative image (scale bar: 100µm) and (C) quantification of IRF7 at the injection site on control versus shIRF7 hemispheres confirming IRF7 knockdown. (D) Representative images (scale bar: 100µm) and (E) quantification of IRF7 within eGFP+ transduces regions in Experiment 4. (F) CIE reduced postsynaptic molecular assessment of E/I ratio in aIC of rats that received AAV8.scCON but not AAV8.shIRF7 injections. (G) CIE reduced PSD95 in the aIC of rats that received AAV8.scCON but not AAV8.shIRF7 injections. (H) Representative images of PSD95. Scale bar: 50µm (I) CIE had no effect on GPHN in aIC in Experiment 4. (J) Representative images of GPHN. Scale bar: 50µ
